# Identification of High-Risk Cells in Single-Cell Spatially Resolved Transcriptomics Data Using DEGAS Spatial Smoothing

**DOI:** 10.1101/2025.01.30.635803

**Authors:** Debolina Chatterjee, Justin L. Couetil, Ziyu Liu, Kun Huang, Chao Chen, Jie Zhang, Michael A. Kalwat, Travis S. Johnson

**Affiliations:** Indiana University School Of Medicine, Indianapolis, IN; Purdue University, West Lafayette, IN; Stony Brook University, Stony Brook, NY; Indiana Biosciences Research Institute, Indianapolis, IN

## Abstract

**Summary:** The examination of high-risk cells and regions in tissue samples from spatially resolved transcriptomics platforms offers meaningful insights into specific disease processes. For existing methods, while cell types or clusters can be identified and associated with disease attributes, individual cells are unable to be associated in the same manner.

**Method:** Diag-nostic Evidence GAuge of Single-cells and spatial transcriptomics (DEGAS), solves the above problem by employing latent representations of gene expression data and domain adaptation to transfer disease attributes from patients to individual cells from single-cell RNA sequencing datasets. In this research, we present and evaluate DEGAS’s versatility in adapting to data arising from various single-cell spatially resolved transcriptomics (scSRT) platforms. DEGAS successfully identified high-risk cells and regions in liver hepatocellular carcinoma and skin cutaneous melanoma, which were validated through known markers. Additionally, DEGAS was applied to our newly generated Type II Diabetes Xenium dataset revealing high-risk cells within the tissue samples.

**Availability and Implementation:** The DEGAS software can be accessed at https://github.com/tsteelejohnson91/DEGAS. For the updated smoothing functions and associated codes, visit https://github.com/dchatter04/DEGAS-Spatial-Smoothing. Sources for the datasets reviewed are detailed in their respective sections. A description of some datasets, along with extra tables and figures, is provided in the Supplementary Materials file. Our newly generated Xenium data for Type II Diabetes can be found at https://doi.org/10.7303/syn68699752.

## 1 Introduction

Single-cell spatially resolved transcriptomics (scSRT) merges sequencing and imaging techniques to study tissue samples with high precision, offering crucial insights into disease mechanisms. Over recent years, scSRT data have evolved to include both spatial and gene expression details, along with related histological images of tissue specimens. Some of the popular platforms that provide such datasets are 10X Genomics’ Xenium [8], MERFISH [3], segFISH+ [5] and Nanostring Technologies’ CosMx [7], among others. Although these data give detailed information about single cells, patient-level attributes such as the overall survival or other physiological characteristics can not be directly attributed to individual cells. Furthermore, these datasets are limited by sample size, making it difficult to generalize findings. On the other hand, bulk RNA-seq datasets from platforms such as the Gene Expression Omnibus (GEO) or The Cancer Genome Atlas (TCGA) reveal information on diverse disease characteristics for patients, such as diagnostic details like disease subtype, status, prognostic information like survival, and treatment responses, but they lack the granular single-cell details. This limitation hinders the identification of cell subsets linked to disease characteristics, especially when disease-related cells are mixed with non-disease-related cells. Thus, there is a pressing need for methods to translate information from scSRT data to the patient level. Using the rationale that single-cell RNA-seq (scRNA-seq) data and patient-level transcriptomic data (such as RNA-seq with clinical annotations) share the same gene set and, therefore, a common feature space, there is a natural link allowing efficient information transfer between these data types to uncover associations between patients and cells. Diagnostic Evidence Gauge of Single-cells and spatial transcriptomics (DEGAS) [9] uses transfer learning methods like domain adaptation [10] and multitask learning [2] to link individual cells to disease risk for scRNA-seq data. In this article, we demonstrate how DEGAS was modified to utilize the spatial information from scSRT datasets. In this article, we demonstrate how DEGAS was modified to utilize the spatial information from scSRT datasets and its utility in liver hepatocellular carcinoma (LIHC), skin cutaneous melanoma (SKCM), and Type II Diabetes (T2D).

## 2 Methodology

The DEGAS framework (Fig. 1 A) is designed using Python’s TensorFlow library and provides an interface in R. In this article we skip the detailed explanation of the framework as it can be found in the original publication [9].

**Figure 1.**
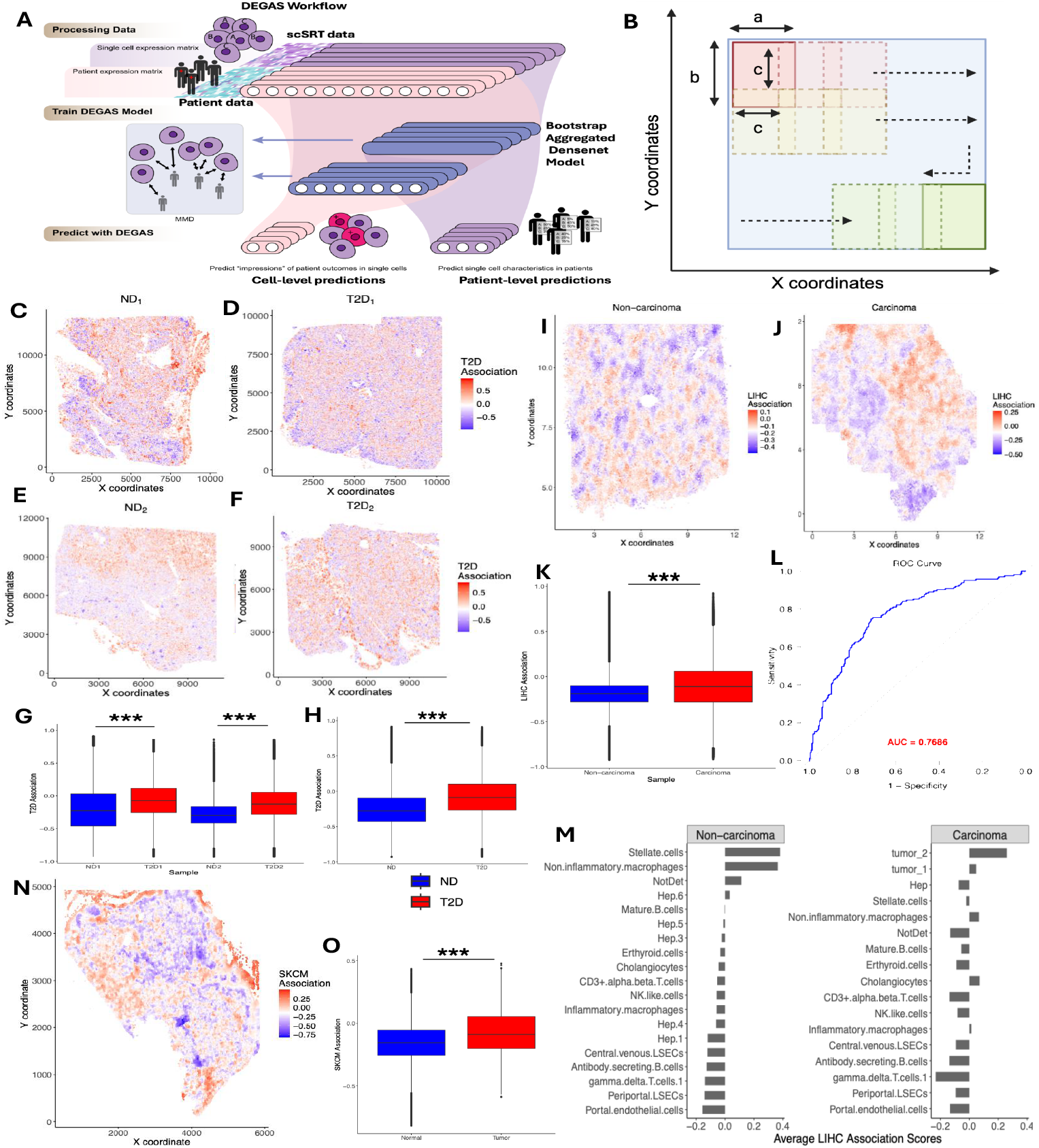
The DEGAS workflow, showing the key inputs and outputs, is overall similar to that of [9]. **B)** Sliding windows progressively shift to capture the entire tissue, enabling localized spatial smoothing of the disease association scores for SwiS and FoVs algorithms. **C, D, E, F)** Spatial visualization of T2D association scores generated by the DEGAS+SWiS model for Xenium tissue samples: *ND*_1_ (panel C), *T* 2*D*_1_ (D), *ND*_2_ (E), and *T* 2*D*_2_ (F), respectively. Legends display the scaled T2D disease association scores. **G)** Boxplot showing a significant difference in T2D association scores across samples, with higher scores observed in T2D samples compared to non-diabetic samples (∗ ∗ ∗ indicates significant p-values). **H)** Combining disease association scores from non-diabetic and T2D samples separately reveals higher overall T2D association scores in diabetic samples. **I, J)** Spatial visualization of LIHC association scores generated by the DEGAS+FoVS model for CosMx tissue samples: Normal (panel I) and LIHC Carcinoma (panel J), respectively. **K)** LIHC association scores for the Carcinoma sample are significantly higher than those of the Non-carcinoma sample. **L)** ROC curve showing patient-level survival prediction by the DEGAS model for classifying 3-year overall survival status in Liver CosMx data. **M)** LIHC association scores by various cell types across the Non-carcinoma and Carcinoma samples, respectively. **N)** Spatial visualization of SKCM association scores generated by the DEGAS+SWiS model for a Xenium tissue sample. **O)** The SKCM association scores for cells within the tumor region are significantly higher than those in the stromal and other non-tumor regions.

The DEGAS model preprocessing and training pipeline is largely the same between scRNA-seq and scSRT data. After training the model (Supplementary Algorithm 1), for each single cell, the patient-level disease association (DA) scores are predicted. Higher DA scores can be interpreted as higher-risk of the disease. The key modification required to adapt to spatial datasets lies in the post-processing steps.

### 2.1 Postprocessing

To adapt to spatial platforms, we use localized smoothing on the predicted DA scores to preserve the spatial context of the tissue and prevent oversmoothing. Let us denote the DA scores, which are scaled within [*−*1, 1] by *S*^scaled^. We define windows based on pixel coordinates from the metadata and locally smooth the DA scores by the Spatial Window Smoothing algorithm below:

#### SWiS Algorithm

Let *𝒞* = *{c*_1_, *c*_2_, …, *c*_*N*_ *}* denote the collection of *N* cells. Each cell *c*_*i*_ is associated with spatial coordinates (*x*_*i*_, *y*_*i*_) *∈* ℝ ^2^, and the DA acores 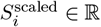, where ℝ is the set of real numbers. Let *𝒟* be set of all cells, where each cell is represented by its spatial coordinates and DA scores, i.e., 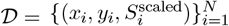, and focus on a small patch of the tissue that spans a 2-dimensional window with 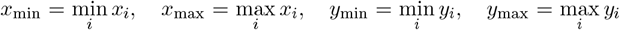 window starting at position (*x, y*) defines a rectangular patch 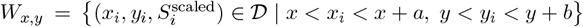, where *a* is window width (X-axis range), *b*: window height (Y-axis range), *c* is step size (stride along X and Y) (as displayed in Fig. 1 B). To ensure the sliding window works well will adequate number of cells (at least *n*_min_), while defining the window, we exclude cells that are too close to the boundary of the window, we use an internal padding *p* to only consider the cells which are in the *core* of it. The *core* of the window exclusing marginal cells is defined as 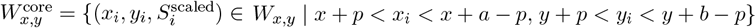.

Starting at the top-left, the window moves rightward until it covers the X-range, then steps down by *c* along the Y-axis, repeating the process until all cells are included. Therefore, the horizontal shift is *x* → *x* + *c* until *x* + *a* > *x*_max_, and vertical shift is *y → y* + *c*, resetting *x* = *x*_min_. This generates a collection of windows 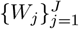, each with a core region 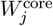.

Within each window 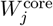 define the set of core cells 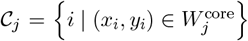. The Algorithm 1 provides a step-by-step description of the SWiS algorithm.

Cells not included in any window can either be excluded from the final output or only their raw scores be considered. A cell can have a list of smoothed DA scores due to overlapping windows and are averaged over all overlapping windows. This ensures full slide coverage while preserving local expression through spatially aware smoothing.

#### FoVS Algorithm

For datasets that are naturally divided into Fields of View (FOVs), instead of defining the sliding window we use the naturally defined FOVs, while other computations remain the same as the SWiS Algorithm 1. Each FOV has its own spatial coordinates and is treated as a discrete region. After smoothing within individual FOVs, the results are combined to reconstruct the full tissue. Next, we present the applicability of DEGAS+SWiS/FoVS in identifying high-risk cells across various diseases and data-types.

## 3 DEGAS identifies high-risk pancreatic cells associated with Type II diabetes using analysis of Xenium data

DEGAS preprocessing steps were conducted on the newly generated Xenium Type II Diabetes (T2D) data (described in Section 2 of the Supplementary material), including filtering for highquality cells and normalizing gene expression to account for technical biases and ensuring robust downstream analysis. For the patient-level bulk RNA-seq data, clinical metadata obtained from the GEO dataset GSE159984 [11], which includes data on 58 non-diabetic (ND) and 27 Type II Diabetic (T2D), was used. This dataset provides important information on patient characteristics such as body mass index (BMI), sex, and T2D status.

There were no cell-type labels in the scSRT data and the bulk RNA-seq data included T2D status classes (T2D vs. ND), we therefore employed the 5x BAg 3-layer *BlankClass* DEGAS model. This model predicted the likelihood of each individual cell being categorized as associated with T2D or ND. We applied SWiS on the DEGAS predicted T2D association scores, with width and height 1000, stride 500, padding 100, 50 minimum cells, and 5 nearest neighbours.

The spatial plot of the DEGAS T2D association scores highlighted individual cells as low T2D-association (blue) or high T2D-association (red) (Fig. 1 C-F). The Wilcoxon rank-sum test revealed that the average T2D-association scores for T2D samples were significantly higher than those of the ND samples (*P* < 0.001, Fig. 1 G,H). This shows the viability of the DEGAS+SWiS approach. We define low-risk cells as those with predicted T2D association scores below the 25^th^ percentile, and high-risk cells as those with scores above the 75^th^ percentile. A threshold of 0.5 is used for the average log_2_-fold change, and only genes expressed in at least 10% of cells are included in the analysis.

Supplementary Tables 2-3 list genes that are significantly upregulated or downregulated (i.e., Benjamini-Hochberg adjusted P-value *≤* 0.05) in the high-risk regions of the Xenium samples. These tables also report the log_2_-fold changes in expressionof these genes between high-risk, lowrisk, and other cells. Supplementary Figures 1-4 present the corresponding volcano plots. Some of the genes that were found to be significantly up/downregulated in the high-risk cells and their association with T2D have been studied, include *PTPRC* [12], *AMY2A* [15, 1] and *CFTR*[13, 6]. Observing these genes in the high-risk cells identified through our algorithm further demonstrates its effectiveness in identifying T2D risk associated cells.

**Figure 2.**
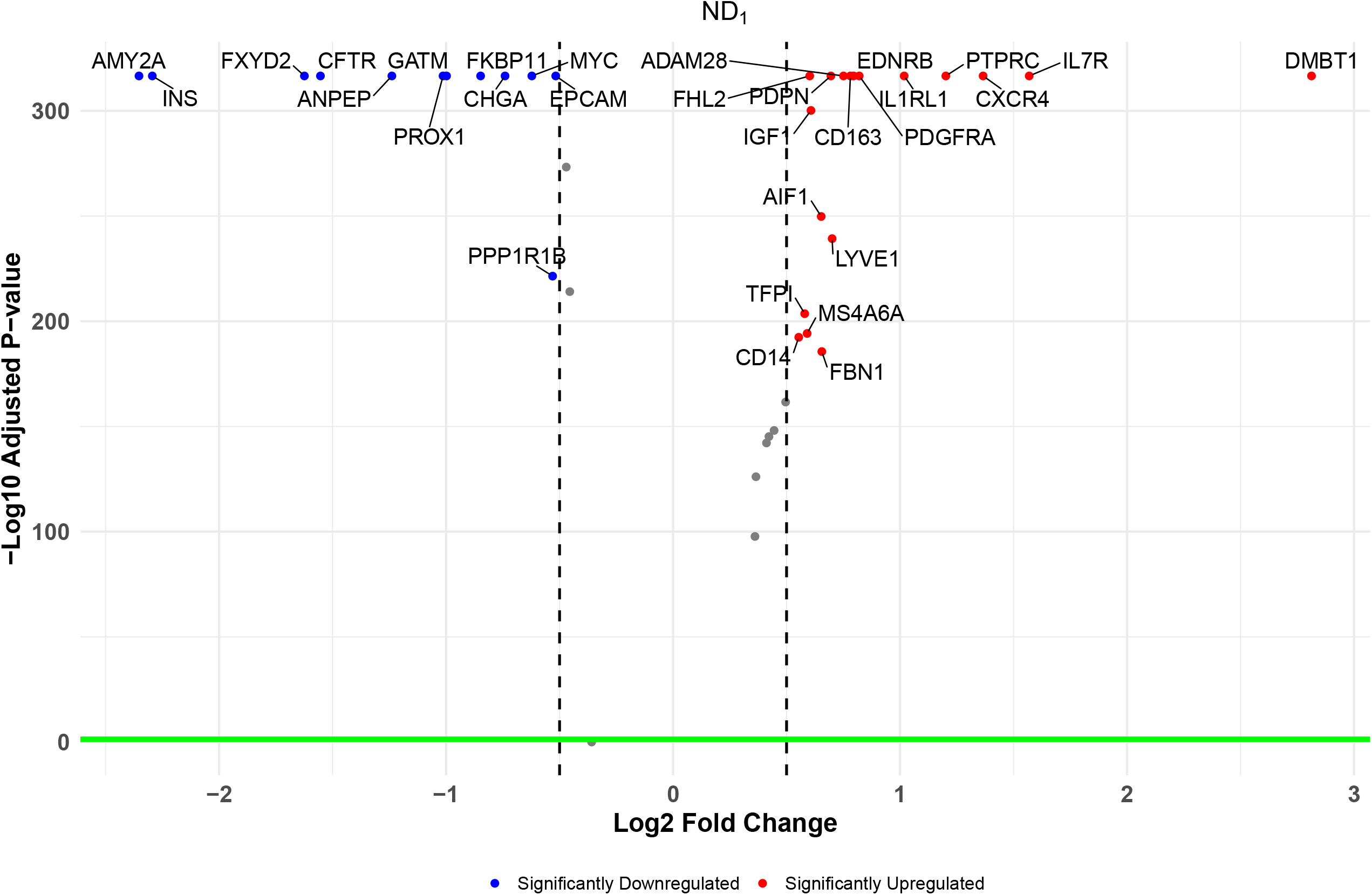

**Figure 3.**
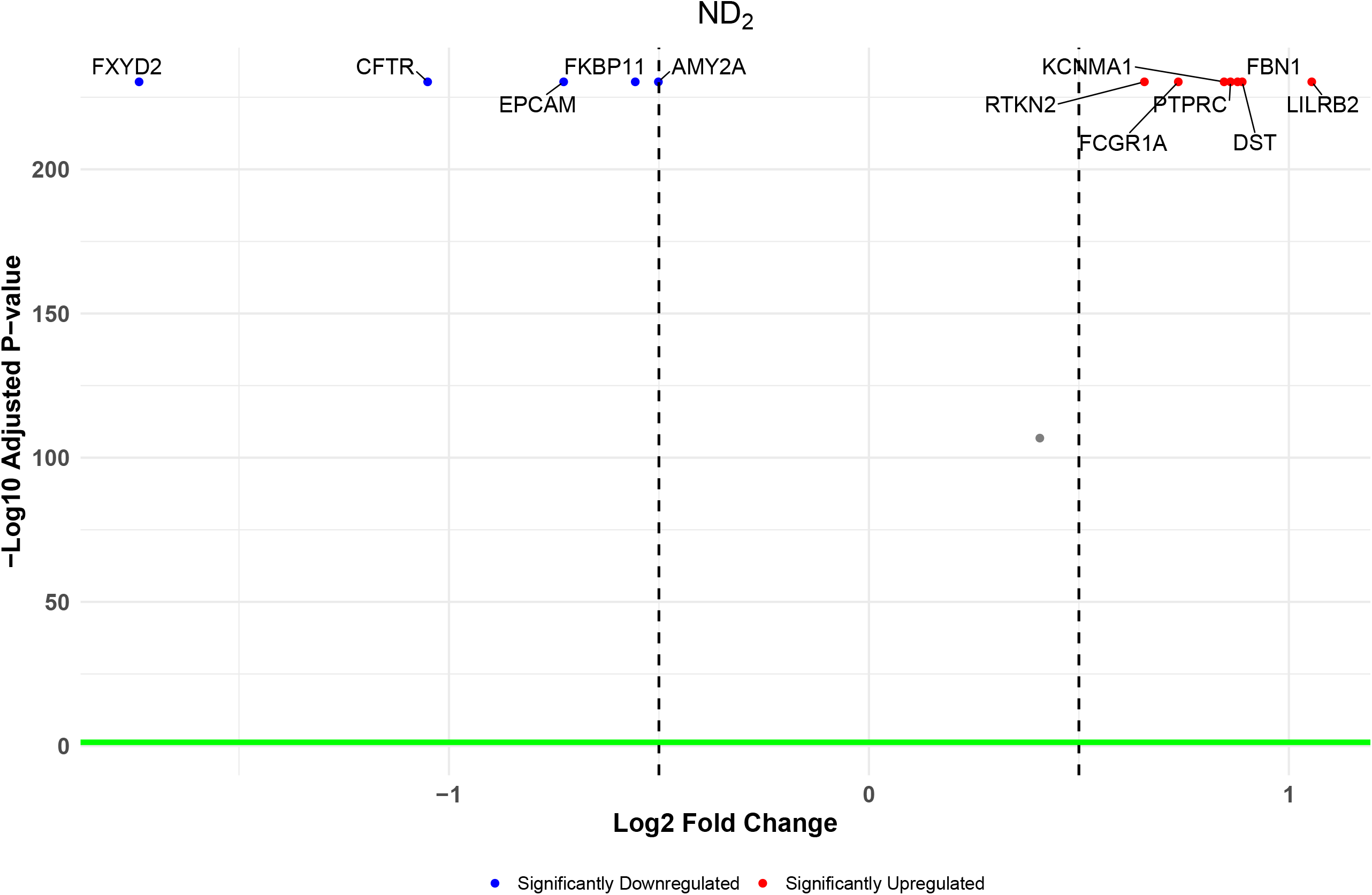

**Figure 4.**
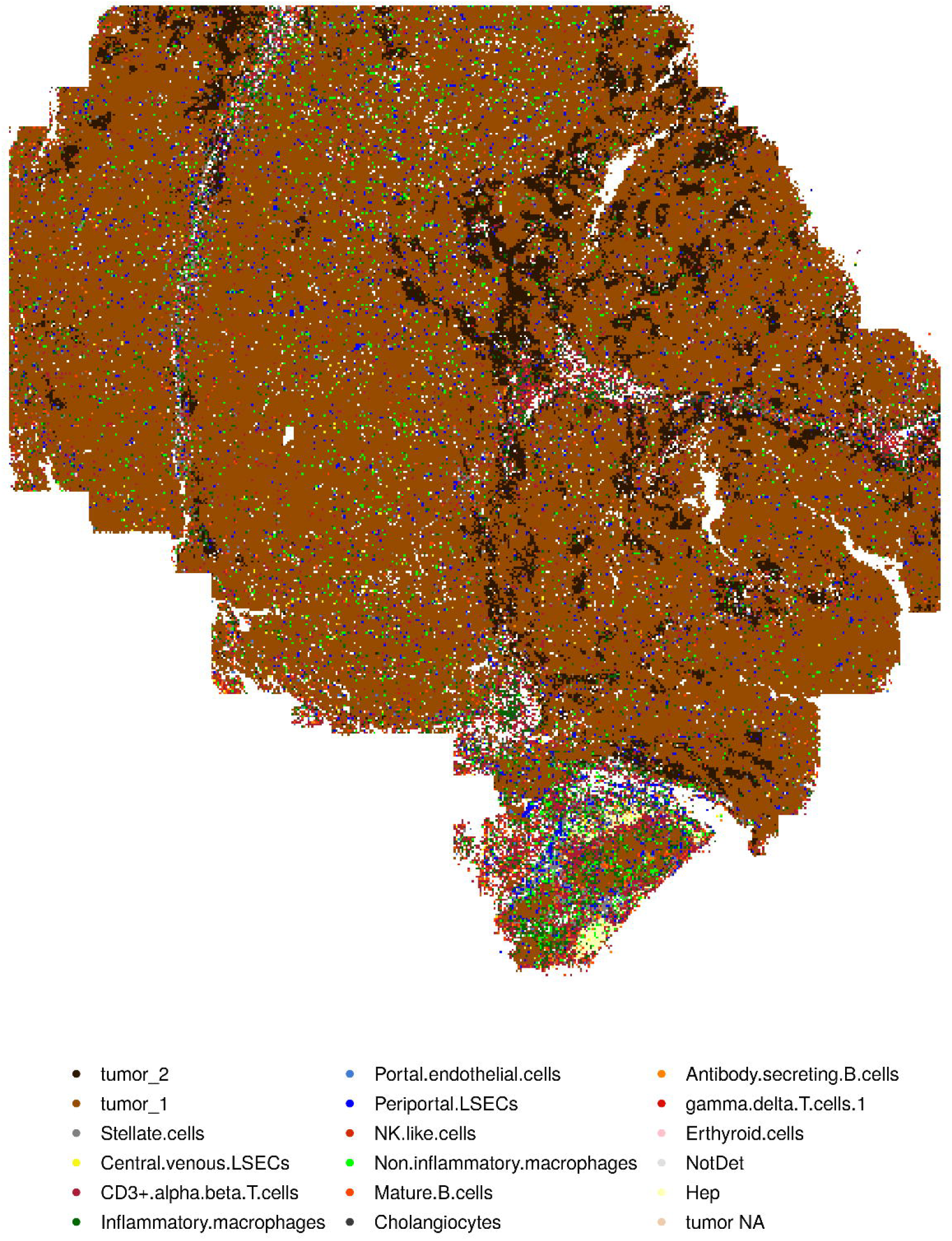

**Figure 5.**
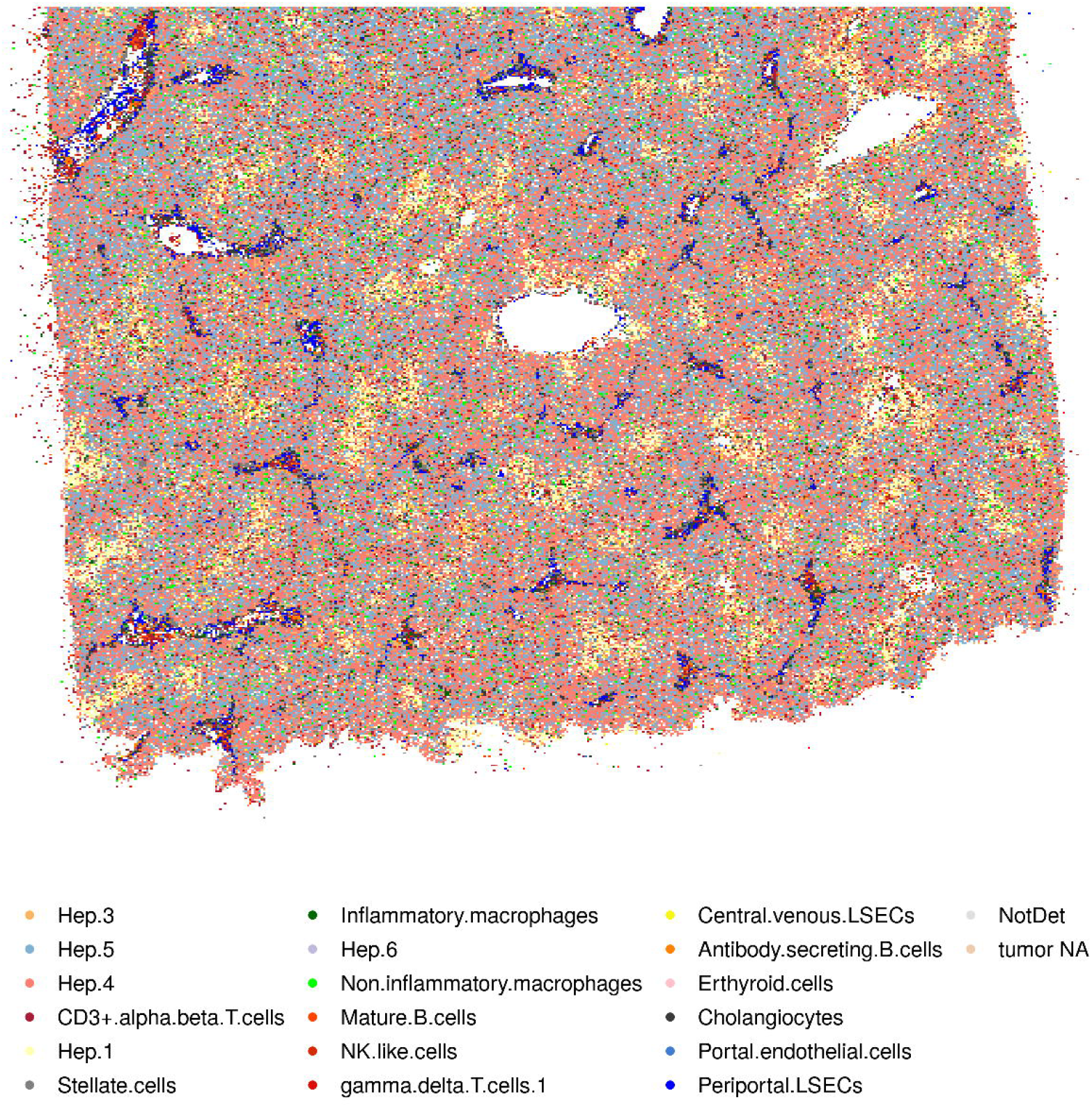

**Figure 6.**
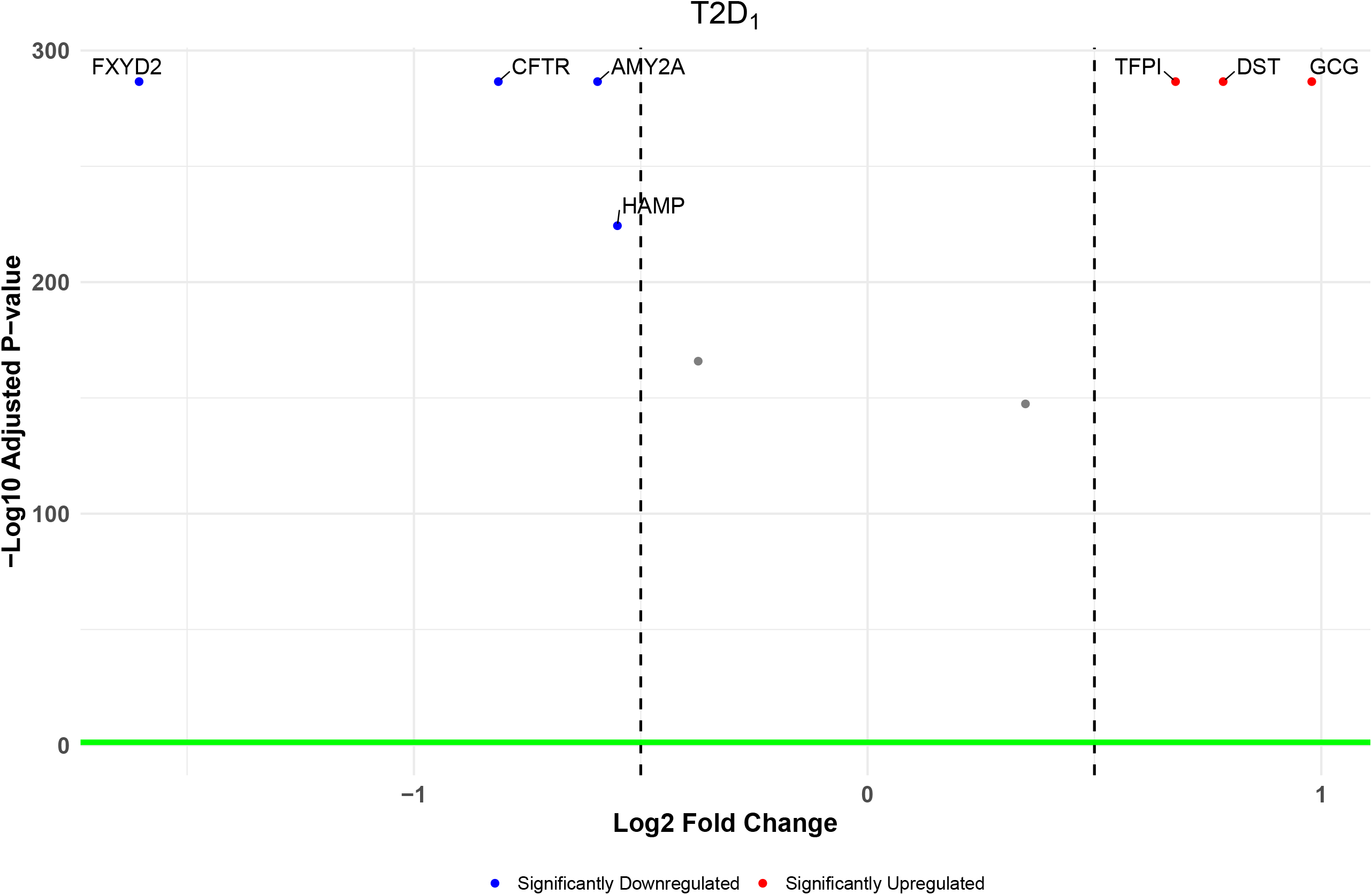

**Figure 7.**
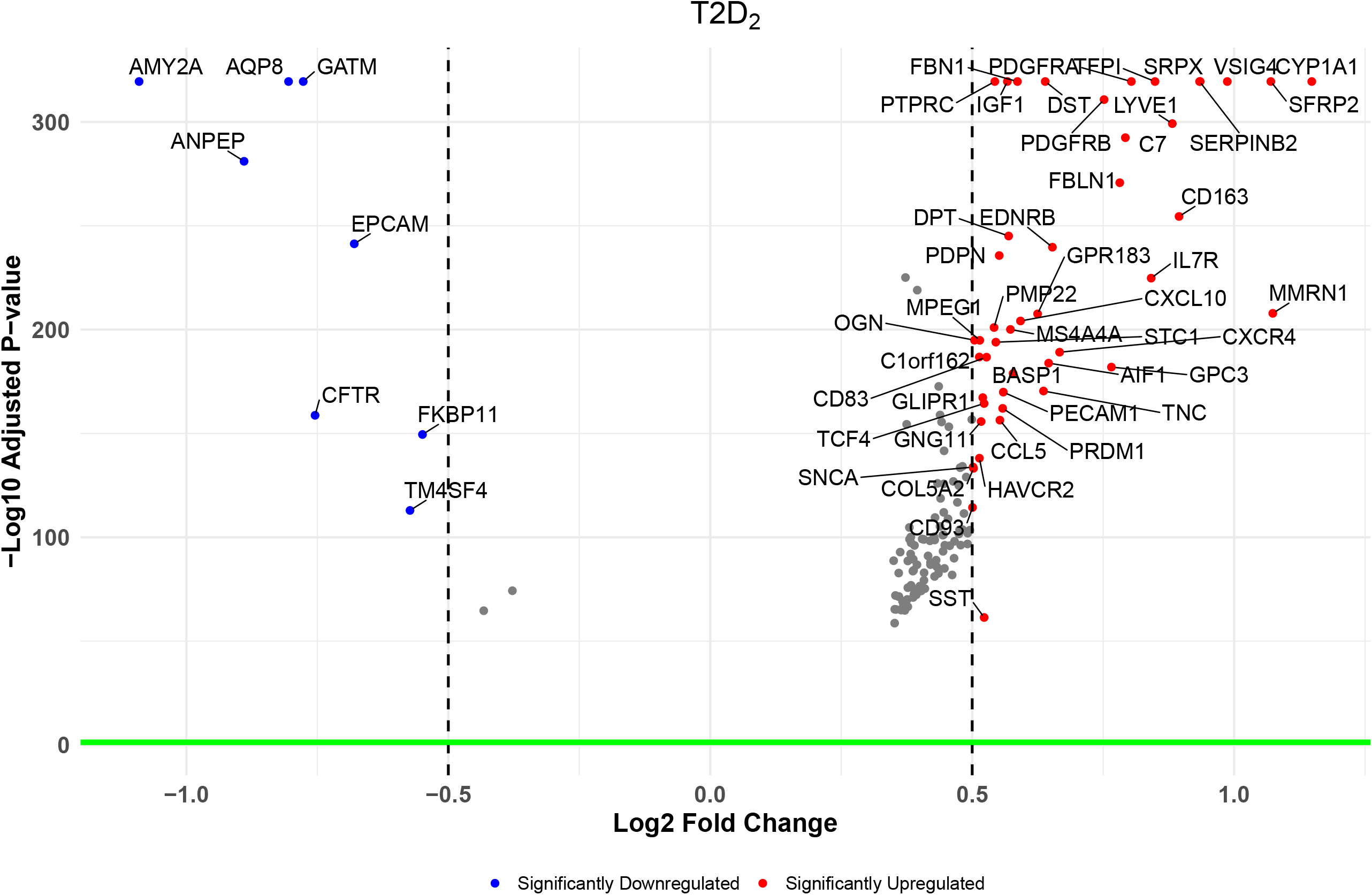

### Algorithm 1 SWiS: Spatial Window Smoothing

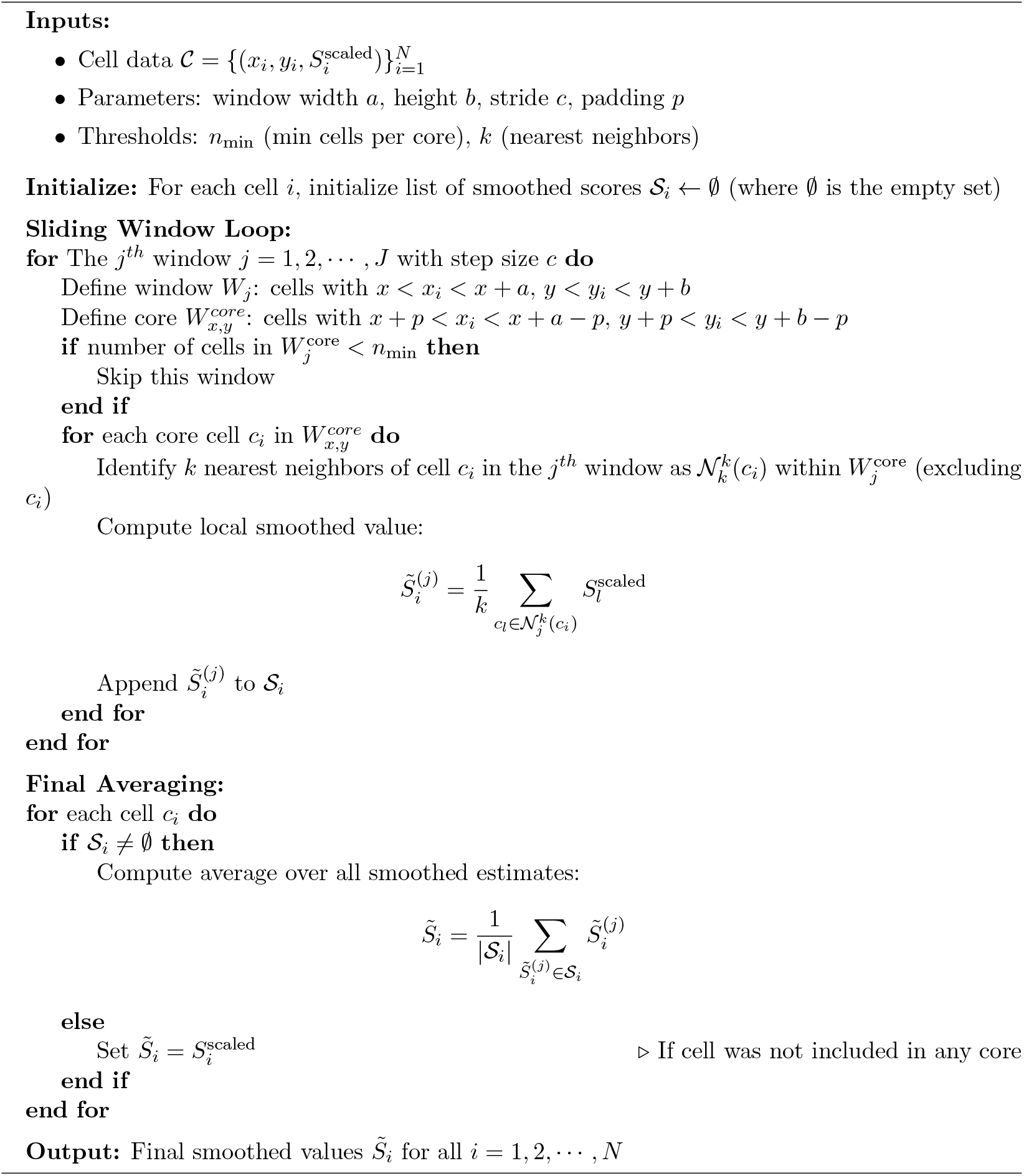

## 4 DEGAS identifies high-risk cell types in CosMx Hepatocellular carcinoma samples

CosMx data is characterized by multiple FOVs that collectively cover the entire tissue area. Within each FOV, individual cells are mapped with precise FOV-specific spatial coordinates and the locations of individual RNA molecules within those cells. We obtained this publicly available Liver Hepatocellular Carcinoma (LIHC) data (see Section 3 of Supplementary material for details). The bulk RNA-seq data with clinical metadata were acquired from the (TCGA LIHC study). This dataset included data from 377 human individuals along with their clinical information, including sex, age, overall survival status, and time.

The gene expression count data for each of the two CosMx samples was individually preprocessed according to DEGAS’s specified preprocessing steps mentioned in the Supplementary Algorithm 1. However, there were no labels distinguishing individual single cells as tumor or normal. Thus, following the identification of a common gene set between the scSRT and the bulk RNA-seq data, the 5x BAg 3-layer *BlankCox* model of DEGAS was utilized. For CosMx data, due to the presence of well-defined FOVs and FOV-specific pixel coordinates, we used the naturally defined FOVs to smooth the LIHC-associations, with 50 nearest neighbours. The spatial map of DEGAS+FoVS LIHC-association scores (Fig. 1 I,J) reveals high LIHC-association regions (red) and low LIHC-association regions (blue) for all cells within a field of view in both the non-carcinoma and LIHC samples. As expected, there is a highly significant difference in the LIHC-association scores between the non-carcinoma and LIHC samples (*P* < 0.001, Fig.1 K). We consider low-risk cells as those with predicted LIHC association scores below the 25^th^ percentile, and high-risk cells as those with scores above the 75^th^ percentile. In the non-LIHC sample, 83219 low- and 83219 high-LIHC association cells were identified, while in the LIHC sample, 115750 low LIHC-association cells and 344691 high LIHC-association cells were identified. This result reveals a significantly higher number of high LIHC-association cells in the cancer sample (*χ*^2^ = 34873.64, *P* < 0.001) compared to the non-carcinoma sample. For patient-level T2D association predictions using the DEGAS model, we utilized binary 3-year survival information from the TCGA dataset. Overall survival time was used as the ground truth, and we excluded censored patients to ensure reliable labeling. Patients who survived beyond 3 years were labeled as 0, while those who experienced an event (death) within 3 years were labeled as 1. The Receivic operating characteristic area under the curve (ROC-AUC) was calculated considering the predicted DEGAS patient level LIHC-association scores. The DEGAS model demonstrated strong discriminatory performance, achieving a high ROC-AUC of 0.7686 (Fig. 1 L).

To further investigate the LIHC-association score for individual cell types, we plotted the overall LIHC-association scores for each cell type within the liver tissues (Fig. 1 M). In the non-carcinoma sample, we observed that stellate cells, and macrophages, representing stromal and immune cells—exhibited elevated scores. In the LIHC (carcinoma) sample, tumor cells, non-inflammatory macrophages, and cholangiocytes showed higher average scores compared to endothelial or gamma delta T cells (Fig. 1 M). Notably, regions in non-carcinoma sample exhibiting high hazard scores for LIHC were enriched with stellate cells, macrophages, mature B cells, and erythroid cells (see Supplementary Figure 5). Macrophages, in particular, are known to contribute to tumor growth by suppressing immune surveillance through the inhibition of cytotoxic T cells [14]. A similar pattern was observed in a DEGAS-based study of prostate cancer, where specific normal tissues were linked to cancer progression [4]. In that study, normal regions with high disease-association scores were found to be infiltrated by macrophages, detectable in both transcriptomic data and histopathological images [4]. In our study, we observed a comparable trend.

## 5 DEGAS identifies high-risk cells in Skin Cutaneous Melanoma samples

For this analysis, the bulk human Skin Cutaneous Melanoma (SKCM) data were acquired from TCGA SKCM study. This reference data contained 293 patients and their overall survival in months. The Xenium scSRT data was sourced from a publicly available 10X Xenium FFPE Human Skin dataset (https://www.10xgenomics.com/) which consists of a single tissue sample. There were no known cell-labels for scSRT data and bulk RNA-seq data has survival times, we applied the *BlankCox* DEGAS model, with overall survival as the patient-level outcome. The SKCM-association scores were calculated, and the SWiS algorithm was applied to find the smoothed DEGAS hazard scores. The SWiS SKCM-association scores were overlaid with the tissue’s spatial coordinates, highlighting areas of higher SKCM association within the tissue (Fig.1 N). Our results demonstrate the effectiveness of DEGAS+SWiS in identifying high-risk regions within Xenium tissue samples (Fig. 1 O).

## 6 Summary

This article demonstrated DEGAS’s seamless application across multiple scSRT platforms and diseases, including 10X Genomics Xenium and Nanostring’s CosMx and various disease types. Additionally, we implemented smoothing techniques, i.e., SWiS and FoVS, that account for spatial localization and created a novel pipeline for scSRT datasets. The application of DEGAS+SWiS/FoVS led to meaningful findings, such as identifying high-risk cell types in liver cancer (LIHC). Many genes associated with Type II Diabetes were found to be significantly upregulated in high-risk cells identified by DEGAS+SWiS. These discoveries position DEGAS with spatial smoothing as a powerful and efficient tool for advancing translational research involving scSRT, offering significant potential for early diagnosis, prevention, and therapeutic interventions.

## Author Contributions

D.C. -Conceptualization, Analysis, Software, Writing; J.L.C., Z.L -Analysis; K.H. -Conceptualization, Review; C.C., J.Z. -Conceptualization, Review, Funding; M.A.K. -Data curation, Review; T.S.J. -Conceptualization, Software, Supervision, Review, Funding.

## Acknowledgement

Sequencing was performed by the Center for Medical Genomics (CMG) at Indiana University School of Medicine (IUSM), which is partially supported by the Indiana University Grand Challenges Precision Health Initiative. This work was supported by the NIH-NCI (R21CA264339) funding received by T.S.J and J.Z., and NIH-NIGMS (1R01GM148970) received by T.S.J., J.Z., and C.C.. General funding support from the IU Precision Health Initiative and AnalytixIN are also acknowledged.

