## Supplementary Material for "Identification of High-Risk Cells in Single-Cell Spatially Resolved Transcriptomics Data Using DEGAS Spatial Smoothing"

**Abstract**

This supplementary document contains detailed description of some datasets, the relevant Algorithm, Tables, and Figures for the manuscript.

### 1 Diagnostic Evidence Gauge of Single-cells (DEGAS) algorithm

Here we briefly describe the main steps in obtained disease association scores through the DEGAS [1] pipeline.

**Algorithm 1** DEGAS Impression Pipeline

**Input:** scSRT data, bulk RNA-seq data with clinical attributes
**Output:** Disease/subtype association scores (*impressions*)
**procedure** DEGAS PIPELINE

**Preprocessing:**

- Align genes (intersection)
- Feature selection (e.g., least absolute shrinkage and selection operator (LASSO), t-tests)
- Normalize expression data (z-scores), scale to $[0, 1]$

**Model Selection:**

- scSRT and bulk both have class labels **Use:** *ClassClass* model
- only bulk has class labels **Use:** *BlankClass* model
- only scSRT has class labels **Use:** *ClassBlank* model
- bulk has survival data **Use:** *BlankCox* model
- scSRT has class labels and bulk has survival data **Use:** *ClassCox* model

**Model Training:**

- Build multitask deep model (e.g., DenseNet)
- Include losses: Cox (survival), classification (attributes), MMD (alignment)
- Perform bootstrap aggregation (BAg) $m$ times
- Tune hyperparameters (batch sizes, dropout, regularization)

**Disease Attribute Mapping:**

- Transfer patient-level attributes to single-cell data
- Generate disease association scores, i.e., impressions

**end procedure**

### 2 Description of dataset used for the Type II Diabetes analysis

For the analysis in Section 3 of the main manuscript, de-identified formalin-fixed parafin-embedded (FFPE) human pancreas tissue was obtained through the National Disease Research Interchange (NDRI) (Table 1). FFPE Tissue blocks were processed into 5 $\mu$m sections, and sections from two donors, two Non-diabetic (ND) and two type II Diabetic (T2D), were mounted directly within the bounding box of the Xenium slide by the Indiana University School of Medicine (IUSM) Histology Lab Service Core. Slides were processed according to the Xenium v1 workflow for in-situ Gene Expression (CG000582) using the Human Multi-tissue and Cancer panel (377 genes) by the IUSM Center for Medical Genomics (CMG) according to the instructions from 10X Genomics Xenium protocol (https://www.10xgenomics.com/). Data were processed and visualized in the Xenium Explorer software. The Xenium data for the four tissue samples can be found at

Table 1: Details of the Xenium Type II diabetes tissue samples. ICH stands for intracerebral hemorrhage

| Sample ID | Gender | Race | Age | BMI | Diagnosis | Cause of Death | T2D Duration |
|---|---|---|---|---|---|---|---|
| $ND_1$ | Male | White | 82 | 28.4 | Normal | Stroke, ICH | NA |
| $T2D_1$ | Male | White | 82 | 37.8 | T2D | Respiratory arrest | 30 yrs |
| $T2D_2$ | Male | White | 77 | 26.3 | T2D | Stroke/ICH | 15 yrs |
| $ND_2$ | Female | White | 78 | 23.7 | Normal | Stroke, ICH | NA |

https://doi.org/10.7303/syn68699752.

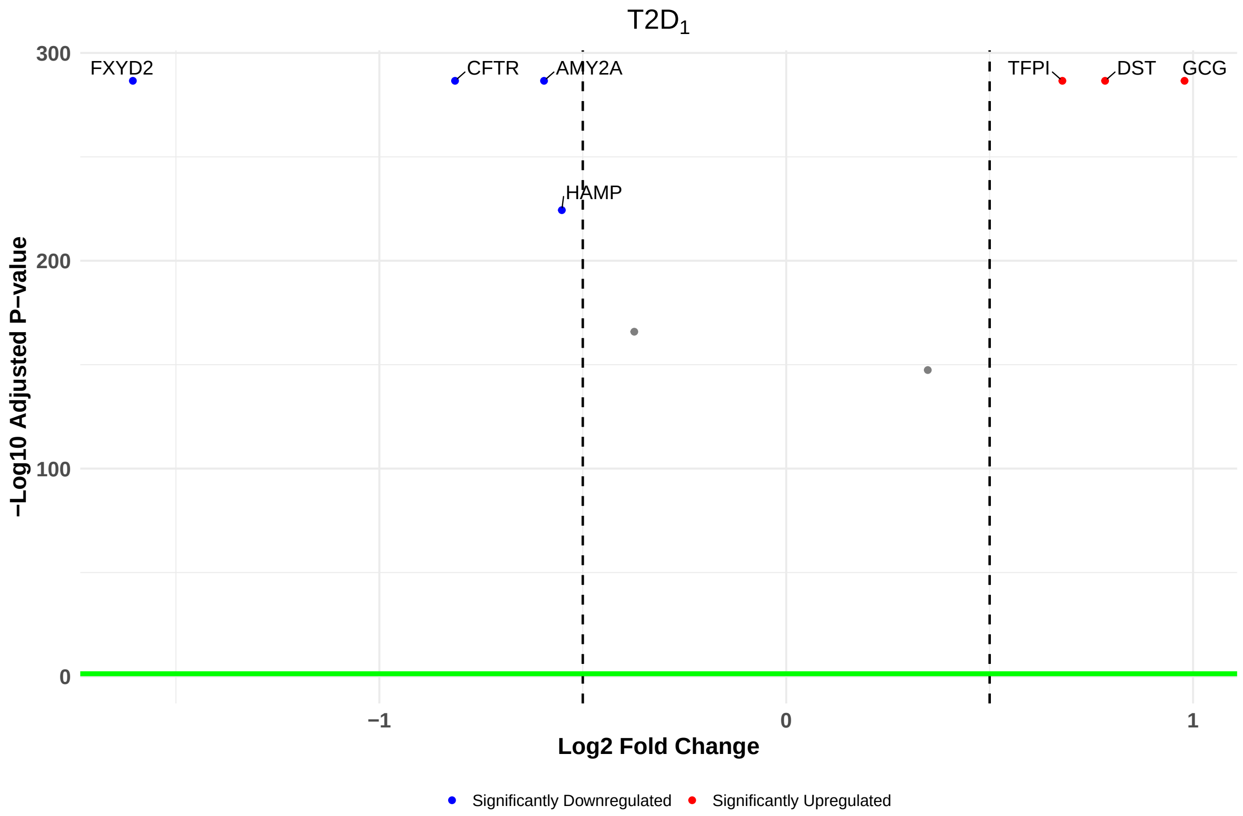


Figure 1: Volcano plots showing significantly upregulated (red) and downregulated (blue) genes considering Benjamini-Hochberg adjusted p-values $\leq 0.05$ and threshold for the average log2 fold-changes in the expression of the genes appearing in at least 10% of cells in the Type II diabetic sample $T2D_1$.

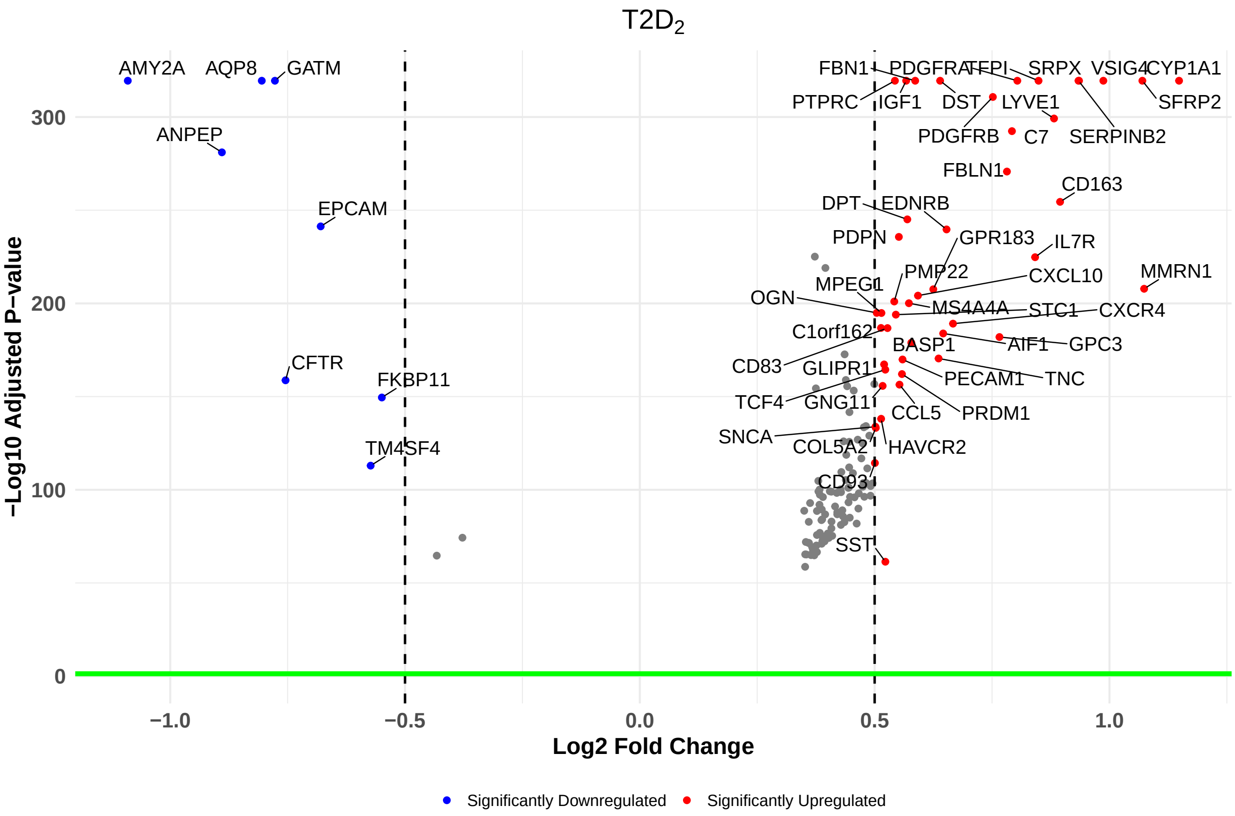


Figure 2: Volcano plots showing significantly upregulated (red) and downregulated (blue) genes considering Benjamini-Hochberg adjusted p-values $\leq$0.05 and threshold for the average log2 fold-changes in the expression of the genes appearing in at least 10% of cells in the Type II diabetic sample $T2D_2$.

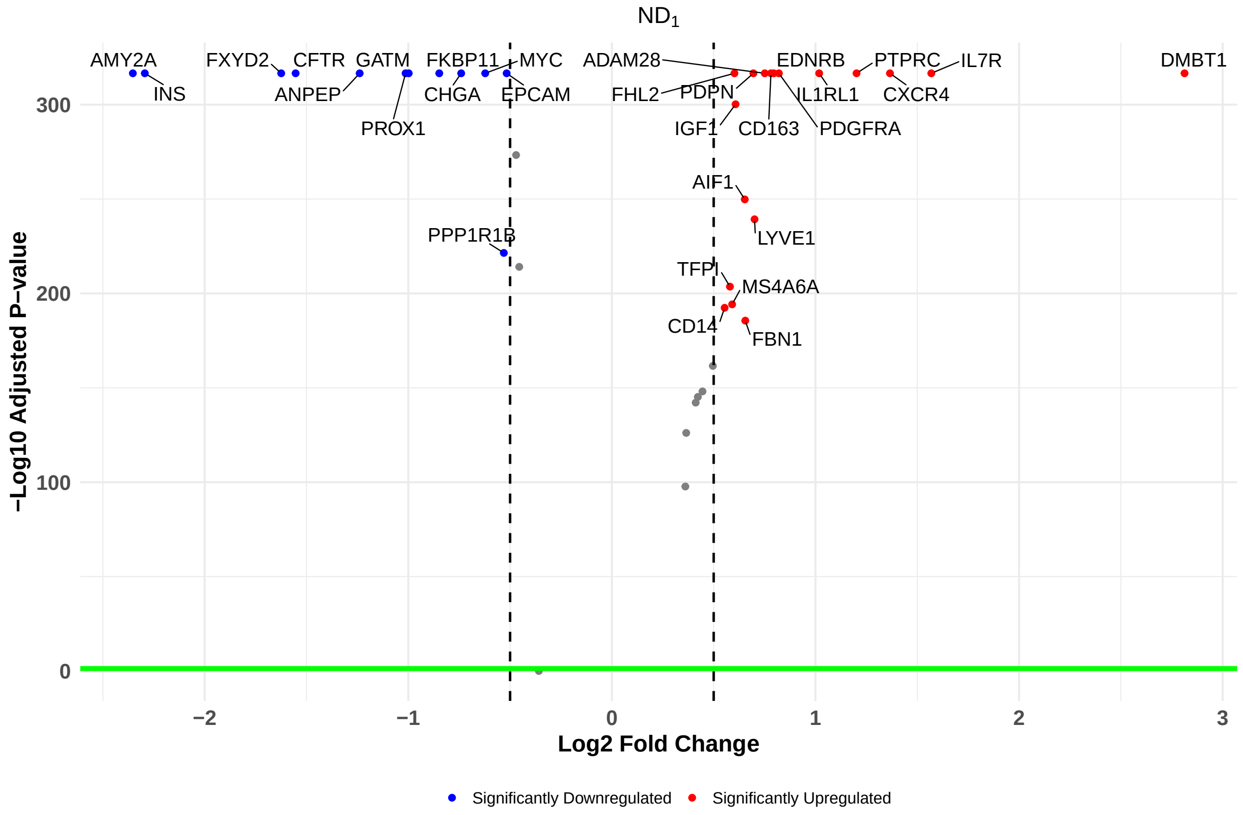


Figure 3: Volcano plots showing significantly upregulated (red) and downregulated (blue) genes considering Benjamini-Hochberg adjusted p-values $\leq 0.05$ and threshold for the average log2 fold-changes in the expression of the genes appearing in at least 10% of cells in the Non-diabetic sample $ND_1$.

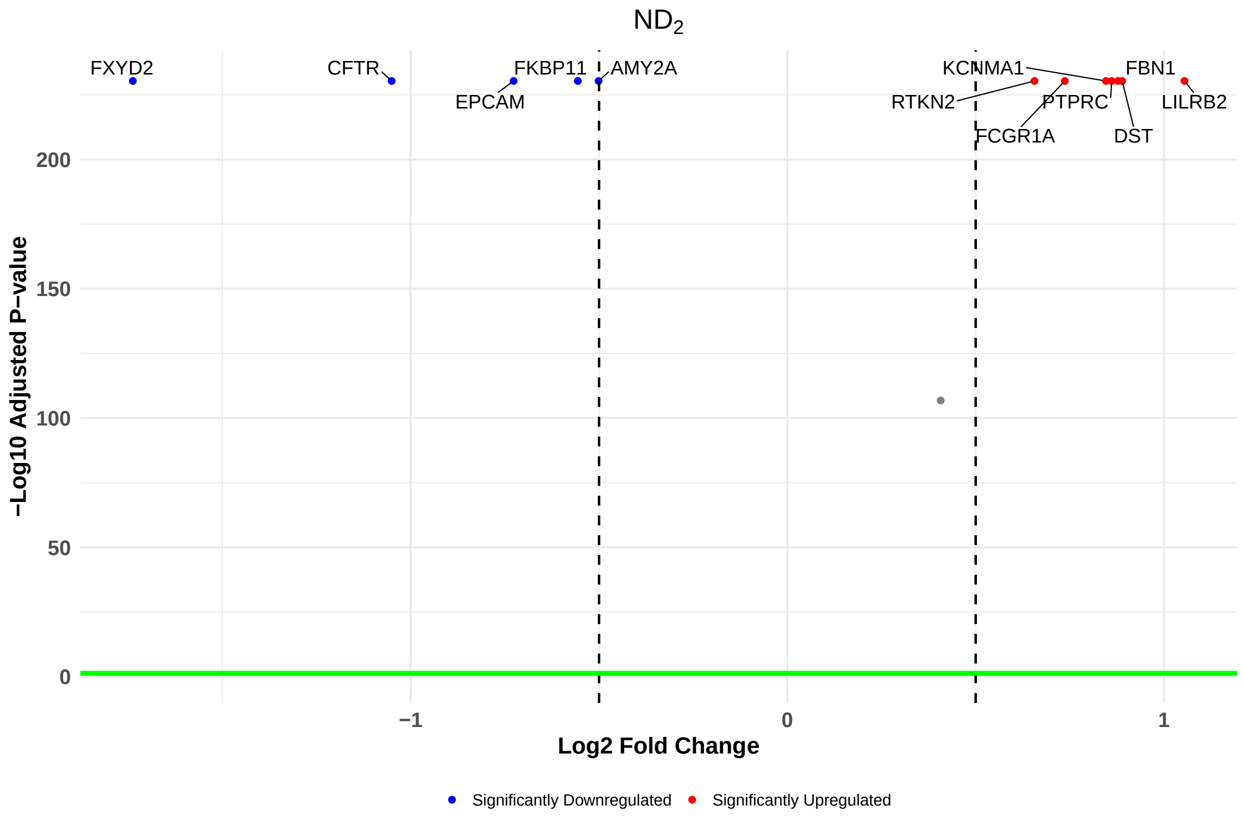


Figure 4: Volcano plots showing significantly upregulated (red) and downregulated (blue) genes considering Benjamini-Hochberg adjusted p-values $\leq 0.05$ and threshold for the average log2 fold-changes in the expression of the genes appearing in at least 10% of cells in the Non-diabetic sample $ND_2$.

Table 2: Upregulated genes with their respective average log 2-fold changes in their expression values in the non diabetic (ND) and Type II diabetic (T2D) conditions.

| **Upregulated genes** | $ND_1$ | $ND_2$ | $T2D_1$ | $T2D_2$ |
|---|---|---|---|---|
| DMBT1 | 2.813 | | | |
| IL7R | 1.569 | | | 0.842 |
| PTPRC | 1.202 | 0.861 | | 0.543 |
| EDNRB | 0.795 | | | 0.653 |
| IL1RL1 | 1.018 | | | |
| CXCR4 | 1.366 | | | 0.667 |
| PDPN | 0.695 | | | 0.552 |
| PDGFRA | 0.82 | | | 0.804 |
| CD163 | 0.781 | | | 0.895 |
| ADAM28 | 0.751 | | | |
| FHL2 | 0.602 | | | |
| IGF1 | 0.607 | | | 0.567 |
| AIF1 | 0.653 | | | 0.646 |
| LYVE1 | 0.701 | | | 0.882 |
| TFPI | 0.58 | | 0.679 | 0.849 |
| MS4A6A | 0.591 | | | |
| CD14 | 0.554 | | | |
| FBN1 | 0.655 | 0.878 | | 0.586 |
| DST | | 0.889 | 0.784 | 0.639 |
| KCNMA1 | | 0.889 | | |
| LILRB2 | | 0.889 | | |
| FCGR1A | | 0.889 | | |
| RTKN2 | | 0.656 | | |
| GCG | | | 0.979 | |
| CYP1A1 | | | | 1.148 |
| VSIG4 | | | | 0.987 |
| SRPX | | | | 0.935 |
| SERPINB2 | | | | 0.934 |
| SFRP2 | | | | 1.07 |
| PDGFRB | | | | 0.752 |
| C7 | | | | 0.792 |
| FBLN1 | | | | 0.782 |
| DPT | | | | 0.57 |
| MMRN1 | | | | 1.074 |
| GPR183 | | | | 0.625 |
| CXCL10 | | | | 0.592 |
| PMP22 | | | | 0.542 |
| MS4A4A | | | | 0.573 |
| OGN | | | | 0.504 |
| MPEG1 | | | | 0.514 |
| STC1 | | | | 0.545 |
| C1orf162 | | | | 0.514 |
| CD83 | | | | 0.527 |

Continued on next page

Table 2 – continued from previous page

| **Upregulated genes** | $ND_1$ | $ND_2$ | $T2D_1$ | $T2D_2$ |
|---|---|---|---|---|
| GPC3 | | | | 0.766 |
| BASP1 | | | | 0.578 |
| TNC | | | | 0.636 |
| PECAM1 | | | | 0.559 |
| GLIPR1 | | | | 0.52 |
| TCF4 | | | | 0.523 |
| PRDM1 | | | | 0.558 |
| CCL5 | | | | 0.553 |
| GNG11 | | | | 0.517 |
| HAVCR2 | | | | 0.514 |
| SNCA | | | | 0.502 |
| COL5A2 | | | | 0.502 |
| CD93 | | | | 0.501 |
| SST | | | | 0.523 |

Table 3: Downregulated genes with their respective average log 2-fold changes in their expression values in the non diabetic (ND) and Type II diabetic (T2D) conditions.

| **Downregulated genes** | $ND_1$ | $ND_2$ | $T2D_1$ | $T2D_2$ |
|---|---|---|---|---|
| AMY2A | -2.353 | -0.501 | -0.595 | -1.09 |
| CFTR | -1.553 | -1.051 | -0.814 | -0.754 |
| EPCAM | -0.517 | -0.727 | | -0.68 |
| INS | -2.294 | | | |
| GATM | -0.998 | | | -0.777 |
| FXYD2 | -1.624 | -1.738 | -1.606 | |
| ANPEP | -1.239 | | | -0.89 |
| PROX1 | -1.013 | | | |
| FKBP11 | -0.848 | -0.556 | | -0.549 |
| MYC | -0.622 | | | |
| CHGA | -0.74 | | | |
| PPP1R1B | -0.531 | | | |
| HAMP | | | -0.551 | |
| AQP8 | | | | -0.805 |
| TM4SF4 | | | | -0.573 |

### 3 Description of the dataset used for Liver Hepatocellular Carcinoma (LIHC) study

This is the description of datasets for the analysis of Nanostring's CosMx data for Liver Hepatocellular Carnimona (LIHC) samples in Section 4 of the main manuscript. The single-cell spatially resolved trabscriptomics (scSRT) dataset was obtained from https://nanostring.com/products/cosmx-spatial-molecular-imager/ffpe-dataset/human-liver-rna-ffpe-dataset/. Figure 5 and 6 respectively show the various cell types whcih were identified in each tissue sample as provided in the publicly available data.

The bulk RNA-seq data with clinical metadata were acquired from the TCGA LIHC study (https://www.cancer.gov/tcga). This bulk RNA-seq dataset included data from 377 human individuals along with their clinical information, including sex, age, overall survival status, and time. is comprised of two FFPE tissue sections analyzed with the Human Universal Cell Characterization Panel (1000-plex). Sample 1 represents normal liver tissue (Male, Caucasian, Age 35), while Sample 2 corresponds to hepatocellular carcinoma tissue (Female, Caucasian, Age 65, Grade G3, Stage II). The gene expression metadata contains individual cell IDs, the FOV pixel coordinates of the centroid of each cell, and the cell type of each individual cell. There are 22 unique cell types. The data contains 340,517 cells in the non-carcinoma sample and 464,126 cells in the LIHC sample.

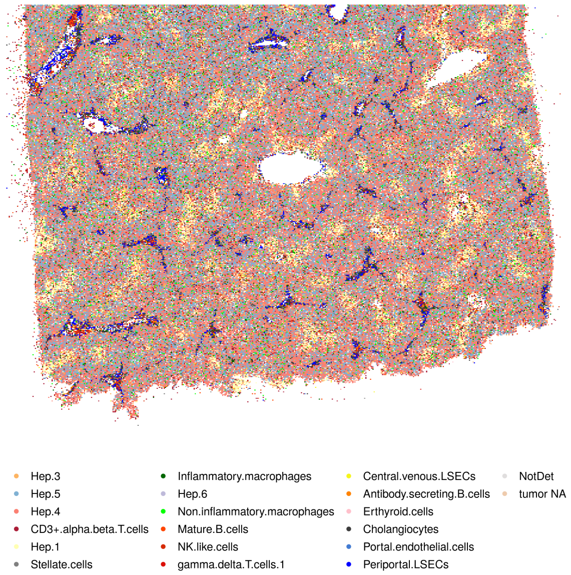


Figure 5: The Non-carcinoma sample showing various cell types. Source: https://nanostring.com/products/cosmx-spatial-molecular-imager/ffpe-dataset/human-liver-rna-ffpe-dataset/

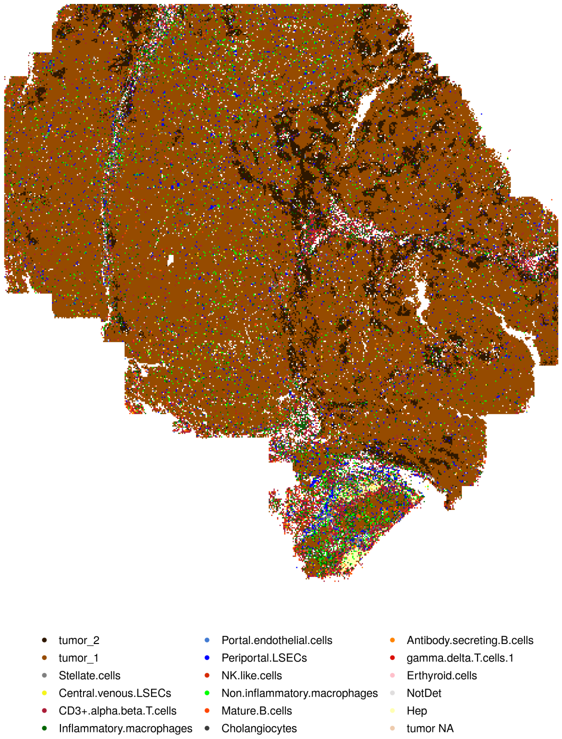


Figure 6: The Liver Hepatocellular Carcinoma (LIHC) sample showing various cell types. Source: https://nanostring.com/products/cosmx-spatial-molecular-imager/ffpe-dataset/human-liver-rna-ffpe-dataset/
